# Epidermal growth factor receptor signaling protects epithelia from morphogenetic instability and tissue damage in *Drosophila*

**DOI:** 10.1101/2022.08.28.505615

**Authors:** Kentaro Yoshida, Shigeo Hayashi

**Author notes:** **Correspondence to:** Shigeo Hayashi, RIKEN Center for Biosystems Dynamics Research, 2-2-3 Minatojima-minamimachi, Chuo-ku Kobe 650-0047, Japan.

## Abstract

Dying cells in the epithelia communicate with neighboring cells to initiate coordinated cell removal to maintain epithelial integrity. Naturally occurring apoptotic cells are mostly extruded basally and engulfed by macrophages. Here, we investigated the role of Epidermal growth factor (EGF) receptor (EGFR) signaling in the maintenance of epithelial homeostasis. In *Drosophila* embryos, epithelial tissues undergoing groove formation preferentially enhanced extracellular signal-regulated kinases (ERK) signaling. In *EGFR* mutant embryos at stage 11, sporadic apical cell extrusion in the head initiates a cascade of apical extrusions of apoptotic and non-apoptotic cells that sweeps the entire ventral body wall. Here, we showed that clustered apoptosis, groove formation, and wounding sensitized *EGFR* mutant epithelia to initiate massive tissue disintegration. We further showed that tissue detachment from the vitelline membrane, which frequently occurs during morphogenetic processes, is a key trigger for the *EGFR* mutant phenotype. These findings indicate that, in addition to cell survival, EGFR plays a role in maintaining epithelial integrity, which is essential for protecting tissues from transient instability caused by morphogenetic movement and damage.

## 1. Introduction

The maintenance of epithelial integrity is essential for animal development and homeostasis. Cell proliferation often produces excess cells that are removed at a later point from the epithelium to maintain optimal cell numbers. The morphogenetic movement of epithelial folding or cell overcrowding applies mechanical stress to the cells to trigger programmed cell death (Eisenhoffer et al., 2012; Levayer et al., 2016; Marinari et al., 2012). These cells are extruded by the contractile activity of healthy neighboring cells. Stabilization of stressed epithelia and rapid repair of epithelial damage due to cell death are especially important in the rapid embryogenesis of *Drosophila* (22 h) compared to zebrafish (72 h) or mice (3 weeks).

*Drosophila* embryogenesis starts from the syncytium, where diploid nuclei fill the surface area yolk occupies the interior. *Drosophila* embryos are internally pressurized and their surfaces supported by the vitelline membrane. Poking a hole in the vitelline membrane causes rapid cytoplasmic leakage; embryos do not survive after vitelline membrane removal. Embryonic morphogenesis proceeds via epithelial folding and invagination in the basal (internal) direction against internal pressure, by increasing the apical surface tension of the epithelium, which drives inward tissue movement (Lecuit and Lenne, 2007). While both apical and basal extrusions have been documented for dying cells in tissue culture and vertebrate tissues (Katoh and Fujita, 2012), *Drosophila* embryos almost always extrude dying cells basally. Here, phagocytic macrophages engulf apoptotic cells so that toxic cell debris is quickly removed and digested for recycling cellular components (Abrams et al., 1993). How the delamination of dying cells is directed toward the interior of the embryo against internal pressure is not understood.

Programmed cell death is induced by various cellular stresses and cell misspecification in mutants of pattern formation genes (Abrams et al., 1993). One major pro-survival system is epidermal growth factor receptor (EGFR) signaling, which regulates extracellular signal-regulated kinases (ERK) signaling in highly dynamic spatiotemporal patterns (Gabay et al., 1997). Massive cell death was observed in EGFR mutants (Price et al., 1989; Schejter and Shilo, 1989). Although EGFR is ubiquitously expressed (Revaitis et al., 2020), the onset of extensive cell death in EGFR mutants is limited to the epidermis of the head and ventral thorax, suggesting that additional factors contribute to the specificity of the prosurvival function of EGFR (Clifford and Schupbach, 1992). In addition, EGFR supports the survival of potentially apoptotic cells in mutants of segmentation genes (Crossman et al., 2018) and promotes the closure of the epithelial opening after apoptotic cell extrusion (Moreno et al., 2019; Valon et al., 2021).

In this study, we addressed the role of EGFR signaling in the maintenance of epithelial integrity in embryos. We showed that EGFR is preferentially activated in the grooves of epithelia, where the apical cell surface is detached from the vitelline membrane. We showed that EGFR and the vitelline membrane, which functions as apical mechanical support, coordinately protect apoptotic epithelial cells from extrusion into the apical side. Here, we discuss the mechanism of maintaining embryonic epithelial tissues from collapsing under mechanical stress from rapid morphogenetic movements.

## 2. Results

### 2.1. EGFR-dependent ERK activity is elevated in the region of epithelial morphogenesis

To reveal the dynamic activation patterns of EGFR during *Drosophila* embryogenesis, we monitored the Förster resonance energy transfer (FRET) probe of ERK activity (Aoki et al., 2017; Komatsu et al., 2011; Ogura et al., 2018). The FRET probe showed repeated stripes of high ERK activity in segmental grooves and the head region, and moderate ERK activity in other epidermal regions of stage 13 and 14 embryos (Figure.1A). The segmental pattern of ERK activity was detected by antibody staining of the activated form (dpERK; Supplementary Figure. S1), as previously reported (Crossman et al., 2018; Gabay et al., 1997; Revaitis et al., 2020). Mutation of *rhomboid (rho)*, which encodes the key protease required for the activation of the major EGFR ligand Spitz (Shilo, 2016), greatly reduced FRET activity (Figure.1B, Supplementary Figure. S1).

**Figure 1.**
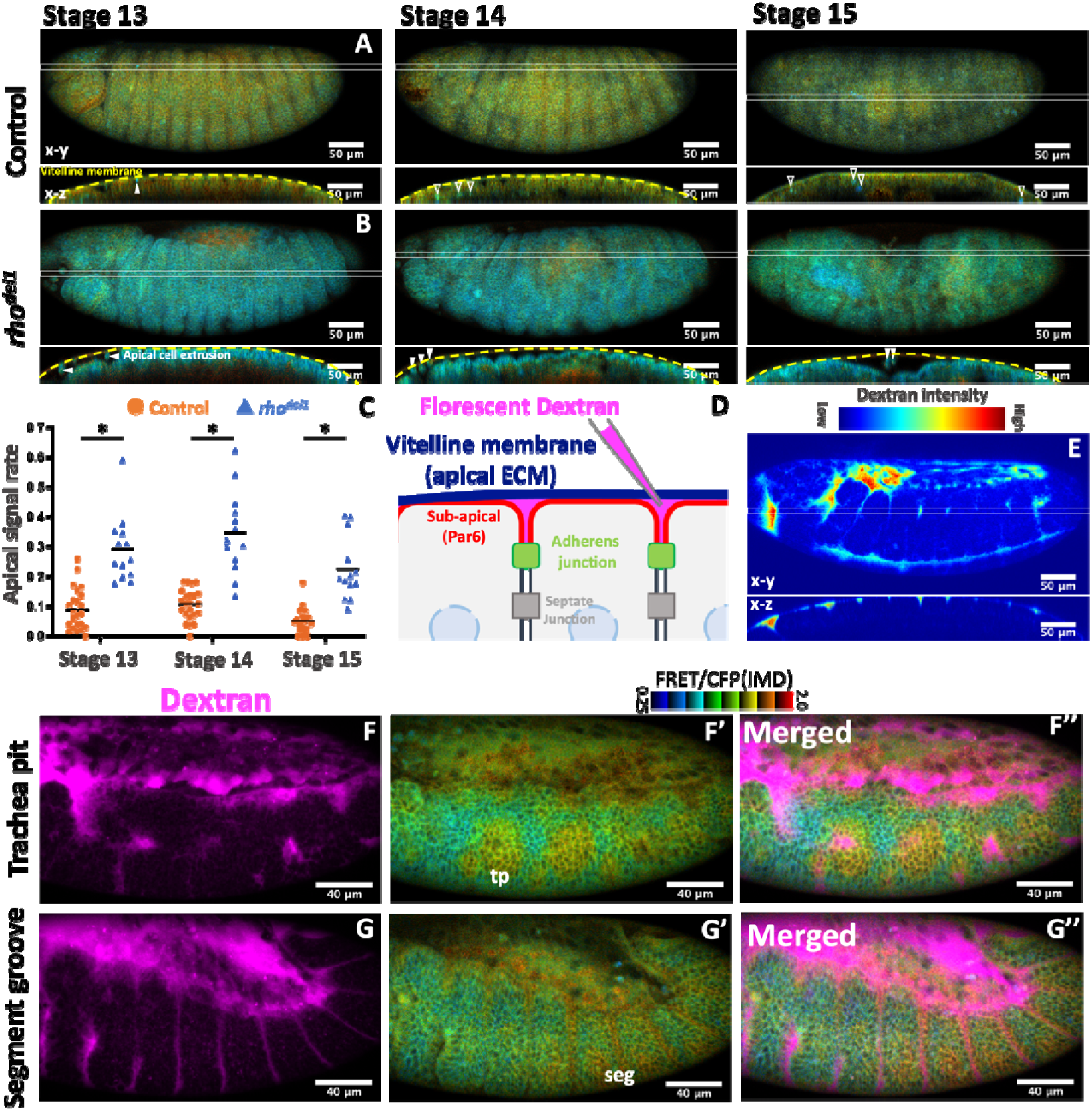
EGFR-dependent ERK activity is elevated in the region of epithelial morphogenesis. **A**. ERK-FRET imaging in a control embryo. The x-y view (z-projected image) and x-z section are shown. The FRET ratio is displayed in IMD format (top right). Open arrowhead, basally localized low ERK-FRET ratio cells; filled arrowhead, apically localized low ERK-FRET ratio cells. **B**. ERK-FRET imaging in a *rhomboid* null mutant embryo. **C**. Quantification of apical cell frequency in control (N = 23) and *rhomboid* null mutant (N = 14) embryos. Asterisk: p < 10^−5^ (Mann–Whitney U test). **D**. Procedure of perivitelline space labeling. **E**. The perivitelline space imaged in stage 11. **F, G**. Simultaneous ERK-FRET and perivitelline space imaging (Supplemental Movie. S1). tp;tracheal pit, seg;segment groove.

In the cross-sectional images, several cells with a low FRET ratio were observed within, or below, the epithelium (Figure. 1A, open arrowhead), with a rare occurrence of the apically localized case (closed arrowhead). Conversely, in *rho* mutant embryos, extruded cells were frequently found in the apical space of the epithelium (Figure.1B, open arrowhead, Figure. 1C).

To further clarify the relationship between EGFR signaling and morphogenetic movement, we performed live imaging of ERK-FRET embryos injected with fluorescent dextran in the space between the vitelline membrane and epidermal cells (perivitelline space, Figure. 1D-G, Supplemental Movie. S1). A high dextran signal was detected in the head-thorax boundary, mouse part (stomodeum), and ventral midline (Figure. 1E). The signal was also detected in the tracheal pit (Figure. 1F) and segmental groove (Figure. 1G). A high level of ERK-FRET activity was detected in the head-thorax boundary, tracheal pit, ventral midline, and segmental boundary (Figure. 1A, 1F,1G, Gabay et al., 1997), where active epithelial folding forms a space between the epithelia and the vitelline membrane. In addition, dextran labeling revealed an extensive space above the amnioserosa, where a high FRET ratio was detected (Figure. 1D-G). Although the high FRET ratio in the amnioserosa was not reduced in *rho* mutants (Figure. 1B), the disintegration of amnioserosa was observed in EFGR mutants (Nishimura et al., 2007; Shen et al., 2013).

These results indicate that EGFR signaling is activated in epithelial tissues undergoing epithelial morphogenesis.

### 2.2. Apical cell extrusion and epithelial collapse in EGFR mutant embryos

The terminal phenotype of EGFR mutants includes massive cell death and loss of a large part of the ventral cuticles, suggesting that the bulk of epidermal cells failed to secrete the cuticle or were lost (Clifford and Schupbach, 1989; Price et al., 1989; Schejter and Shilo, 1989). Observation of all stages up to stage 11 revealed that the bulk shape of the embryo was maintained, however, various local phenotypes existed, such as defects in ventral epidermal patterning and tracheal invagination (Nishimura et al., 2007; Raz and Shilo, 1993).

To understand the process of the massive loss of epidermal cells in EGFR mutants, we performed time-lapse imaging of EGFR mutant embryos. Ventral views of the control embryos showed segment groove formation (t = 2 h) and head involution (t = 4 h) (Figure. 2A). EGFR mutants at early stage 11 (t = 0) showed normal epithelial integrity, as shown by the junctional localization of Par6-GFP and the clustering of salivary gland placode cells (Figure. 2B, orange arrows). At the germ band retraction stage, several condensed cells lacking junctional Par6-GFP appeared in the head region (Figure. 2B; t = 2 h, white arrow), and this phenotype expanded posteriorly. Four hours later, the bulk of the ventral epidermis (ca. 70 %) lost junctional Par6-GFP (Figure. 2B, t = 4h, Figure. 2C).

**Figure 2.**
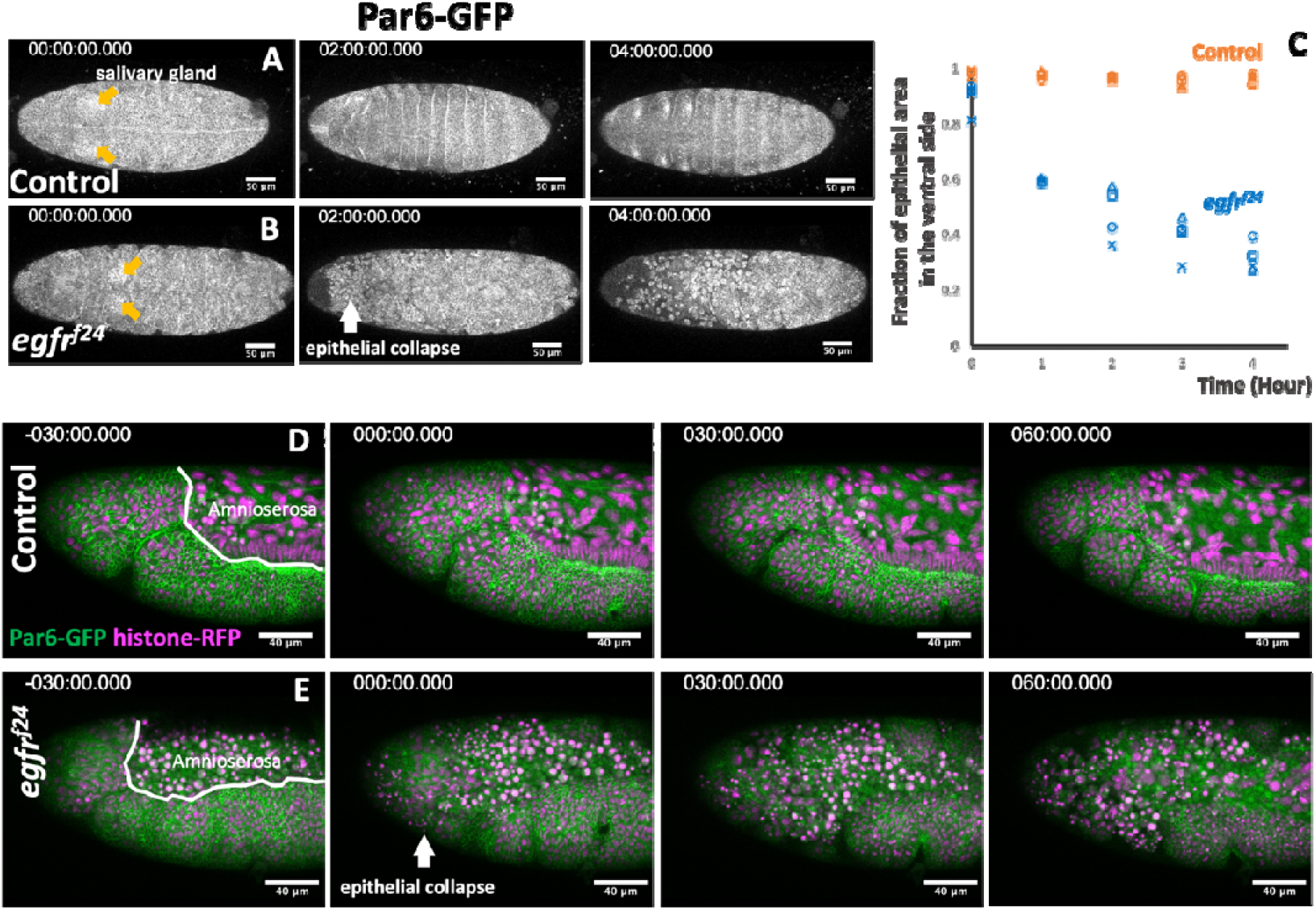
Epithelial collapse in the EGFR mutant embryos. **A, B**. Time-lapse imaging of ventral views of control (A) and EGFR-mutant (B) embryos carrying the junction marker Par6-GFP. Time (t) = 0 was set at the time of salivary gland invagination (orange arrow). The site of the initial epithelial collapse in the EGFR mutant is marked with a white arrow (t = 2 h). **C**. Time course of loss of junctional Par6-GFP localization. **D**. Lateral view of the head of a control embryo. **E**. An EGFR mutant embryo. Amnioserosa had already disintegrated at t = -30 min, followed by the loss of epithelial integrity at t = 0 min. At t = 60 min, the entire head region collapsed.

Time-lapse imaging performed in a lateral view (Supplemental Movie. S2) showed that the amnioserosa was the first to lose continuity in EGFR mutants (Figure. 2D, E. t = -30 min, Supplemental Movie. S2) (Shen et al., 2013). After the onset of tracheal invagination at stage 11, rounded cells were frequently observed on the apical side of the epidermis (Figure. 2E, t = 0, 30 min). Subsequently, the entire head ectoderm of the EGFR mutants lost epithelial connections and collapsed (Figure. 2E, t = 60 min, arrows), whereas the thoracic ectoderm retained its epithelial state.

### 2.3. Apical cell extrusion from apoptotic cell clusters in EGFR mutants

Apoptosis in the head is developmentally controlled by the head patterning hox gene *deformed* (*dfd*), which stimulates the expression of the proapoptotic gene *reaper* (Lohmann et al., 2002; Nassif et al., 1998). Apoptotic cells labeled with the cleaved *Drosophila* caspase 1 antibody (cDcp1) were not detected in the control and EGFR mutant embryos at stage 9 (Figure. 3A, C). In control embryos, the number of apoptotic cells was increased and formed two prominent clusters at the ocular segment in stage 12 (Figure 3B). Those cells were present in the plain of the ectoderm or on the basal side (Figure 3B’, open arrowheads). In EGFR mutant embryos, cDcp1 positive cells increased more extensively. Massive cDcp1 positive cells occupied the apical area of grooves corresponding to the ocular segment, stomodeum, and salivary gland (Figure 3D, closed arrowheads). The clusters of cDcp1 positive cells were also detected in the plain of the ectoderm or on the basal side (Figure 3D’, open arrowheads).

**Figure 3.**
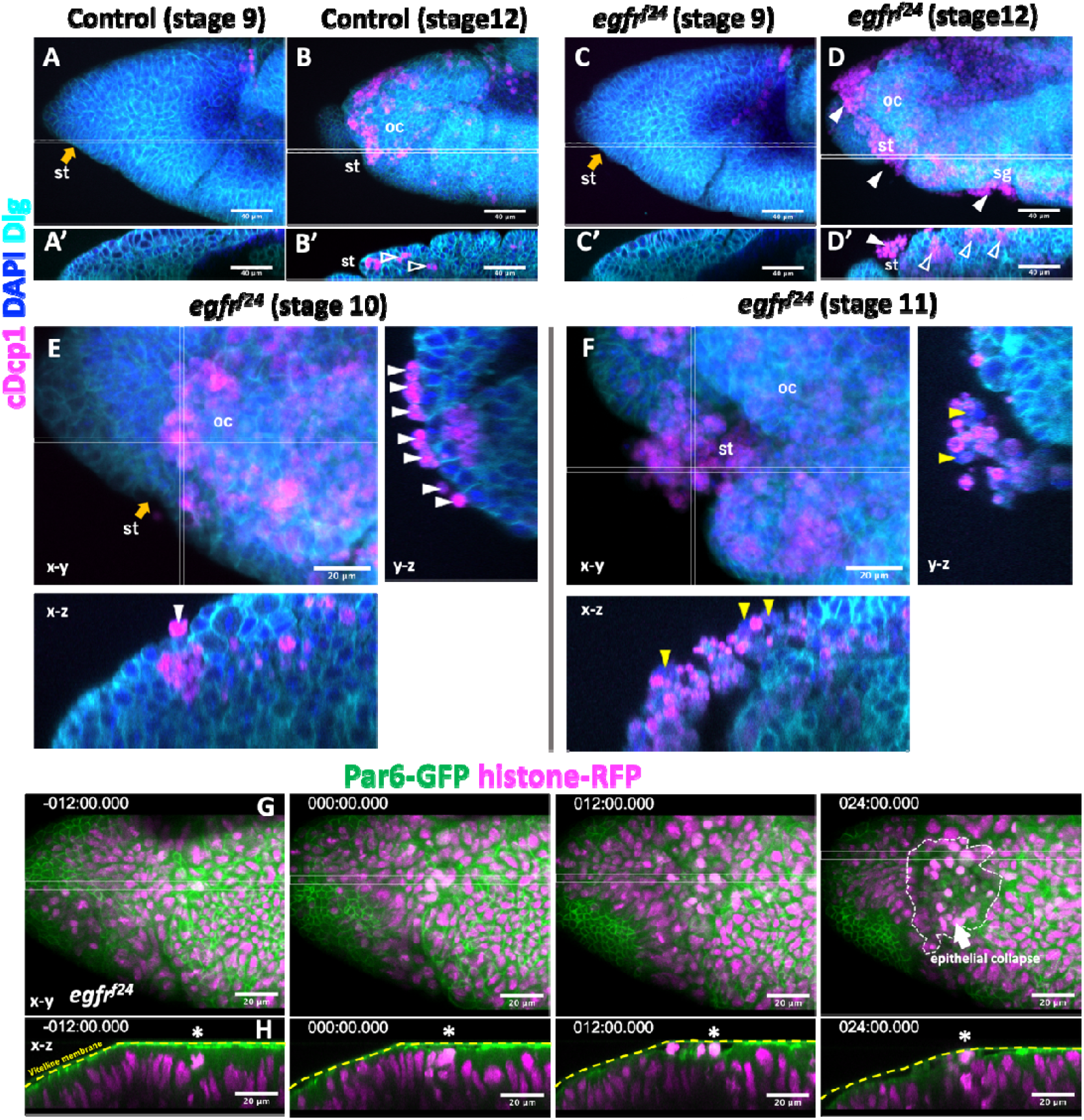
Apical extrusion of apoptotic and non-apoptotic cells in EGFR mutants. **A-F**. Active caspase staining (cDcp1) of control (**A, B**) and EGFR mutant (**C-F**) embryos. In stage 9, no apoptotic (cDcp1 positive) cells were observed within the head epidermis of control and EGFR mutant embryo (A,C). In stage 12, cDcp1 positive cell appeared as clusters within, or basal to, the plane of the epithelium (open arrowheads) of control embryos (**B**). In EGFR mutants, apoptotic cells appeared basal to the epithelium in stage 12 (**D**), however, most appeared apical to the epithelium in stage 12 (white arrowhead). Apical cells were all cDcp1 positive in early stage 10 (**E**). In late stage 11, cDcp1 negative cells were found in the apical location (yellow arrowhead, **F**). **G**. Live imaging of the head region of the EGFR mutant showing an extensive collapse in the ocular segment of the head (arrow in the panel 24 min). Here, t = 0 was set at the time the first apical cell extrusion event was observed (asterisk). st; stomodeum, oc; ocular segment, sg; salivary gland. Orange arrows indicate the invagination region of ectoderms into the stomodeum.

Observation of the ocular segment of EGFR mutants in early stage 10 showed that apically extruded cells were individually attached to the surface of the epithelium, and these cells were all cDcp1 positive (Figure. 3E). In stage 11 embryos after stomodeum invagination, clusters of extruded cells, including both cDcp1 positive and negative cells, were observed (Figure. 3F, yellow arrowheads).

Live imaging of the EGFR mutant heads was performed (Figure. 3G, Supplemental Movie. S3). Before apical cell extrusion, the apical cell surface was tightly juxtaposed to the vitelline membrane (Figure. 3G, t = -12 min). A gap between the apical cell surface and vitelline membrane appeared (Figure. 3G, t = 0). One cell with condensed chromatin was found to move into the apical space (Figure. 3G, asterisk), followed by extrusion of additional cells (white arrowhead), leading to the collapse of the near entire head (Figure. 3G, t = 24 min, Supplemental Movie. S3).

Mitotic cell staining showed that mitosis occurred in clusters within the plains of epithelia in the heads of both control and EGFR mutant embryos (Supplementary Figure. S2A, B). We also observed no mitotic figure in the apically extruded cells of the EGFR mutants (Supplementary Figure. S2B).

### 2.4 Apical cell extrusion at the site of epithelial detachment from the vitelline membrane

To further clarify the relationship between epithelial fold formation and apical cell extrusion, we further investigated posterior segments (Figure. 4A). A ventral view of stage 11 embryos, before the onset of the massive collapse of the ventral epidermis, showed a significantly increased number of apoptotic cells forming clusters in EGFR mutants (Figure. 4B, C). Cross-sectional views of mutant embryos showed that most apoptotic cell clusters were found on the basal side of the epithelia (Figure. 4D, D’, E, E’ open arrowheads). A notable exception was the salivary gland placode, where apoptotic cells formed clusters on the apical side (Figure. 4D, D’, E, E’, orange arrow, closed arrowhead). Time-lapse imaging showed that apical cell extrusion occurred in the salivary gland pit after the invaginating placode formed an open space between the vitelline membrane (Figure. 4F, G, Supplemental Movie. S4). No apical cell extrusion was observed outside the salivary gland placode in the imaged region of mutant embryos.

**Figure 4.**
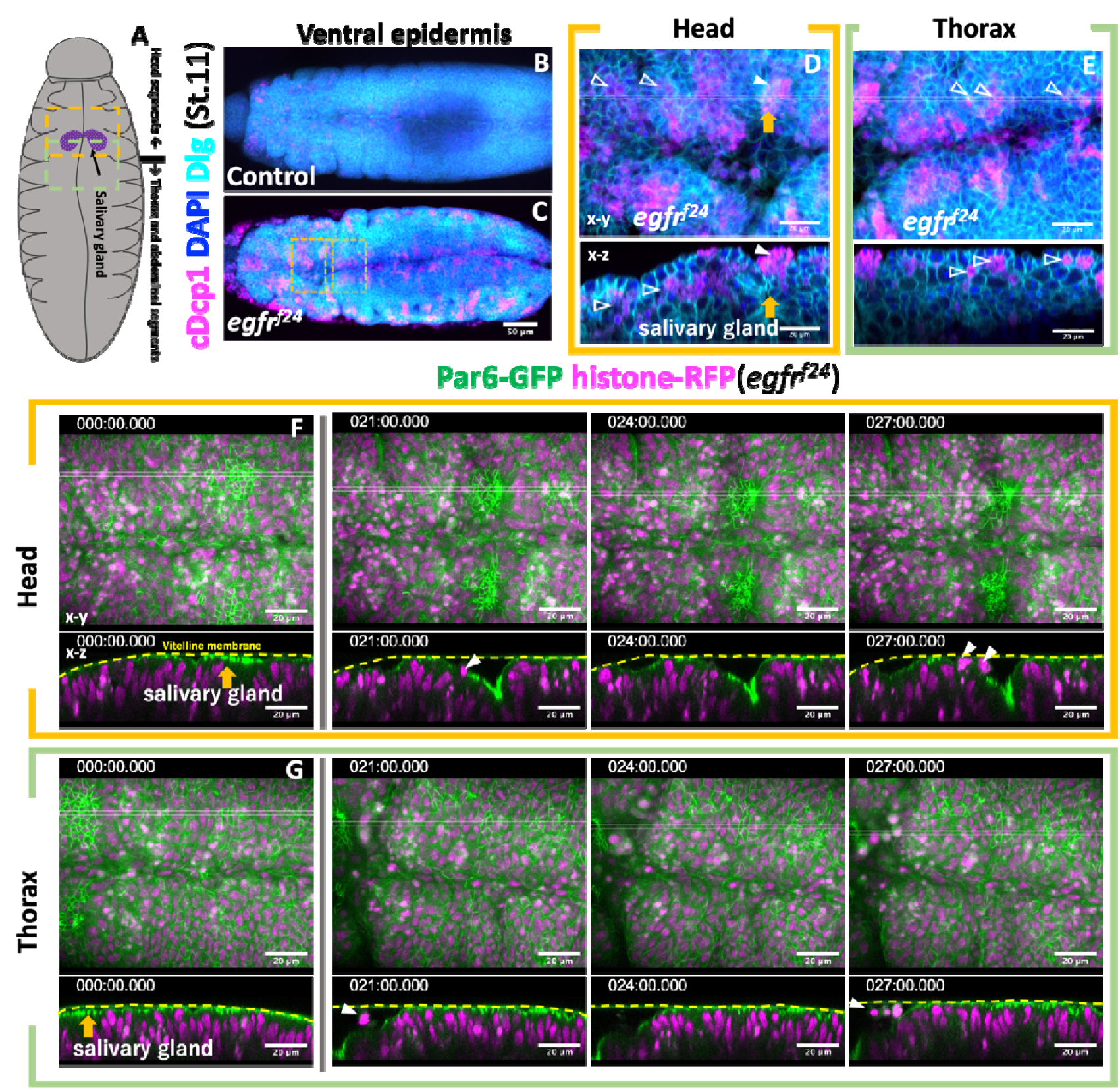
Apical cell extrusion at the site of vitelline membrane detachment. **A**. Schematic of the ventral view of stage 11 embryos. **B-E**. Active caspase staining (cDcp1) of control (**B**) and EGFR mutant (**C**) embryos. Parts of the head and thoracic regions of **C** (dotted box) are enlarged in **D** and **E**. Open and white arrowheads, basally extruded cells; orange arrow, salivary gland invagination. **F, G**. Time-lapse series of the ventral head and thorax imaged from two separate embryos, showing apical cell extrusion in the invaginating salivary gland primordia (orange and white arrowheads). Other parts maintained tight vitelline membrane contact in this stage.

### 2.5 Forced detachment of the vitelline membrane caused the rapid collapse of EGFR mutant epithelia

Considering the results thus far, we reasoned that attachment to the vitelline membrane suppresses epithelial instability under the loss of EGFR function. To test this hypothesis, we observed epithelia mechanically detached from the vitelline membrane.

A stainless-steel needle was used to remove the vitelline membrane from stage 11 embryos, and the resultant embryo fragments were cultured in a petri dish (Figure. 5A-C). EGFR+ tissue fragments expressing the caspase activity reporter Apoliner (Figure. 5D, Bardet et al., 2008) showed a gradual increase in the signal at ca. 4 h, but their tissue structure mostly remained intact over 7 h (N = 13 cultured fragments, Figure. 5E). Strikingly, when EGFR mutant epithelial tissues were cultured, they rapidly lost integrity and nearly completely collapsed within 5 h (N = 13 cultured fragments, Figure. 5F, Supplemental Movie. S5). Cross-sectional views showed that EGFR mutant epidermal fragments were folded and extruded several cells apically before completely collapsing (Figure. 5G).

**Figure 5.**
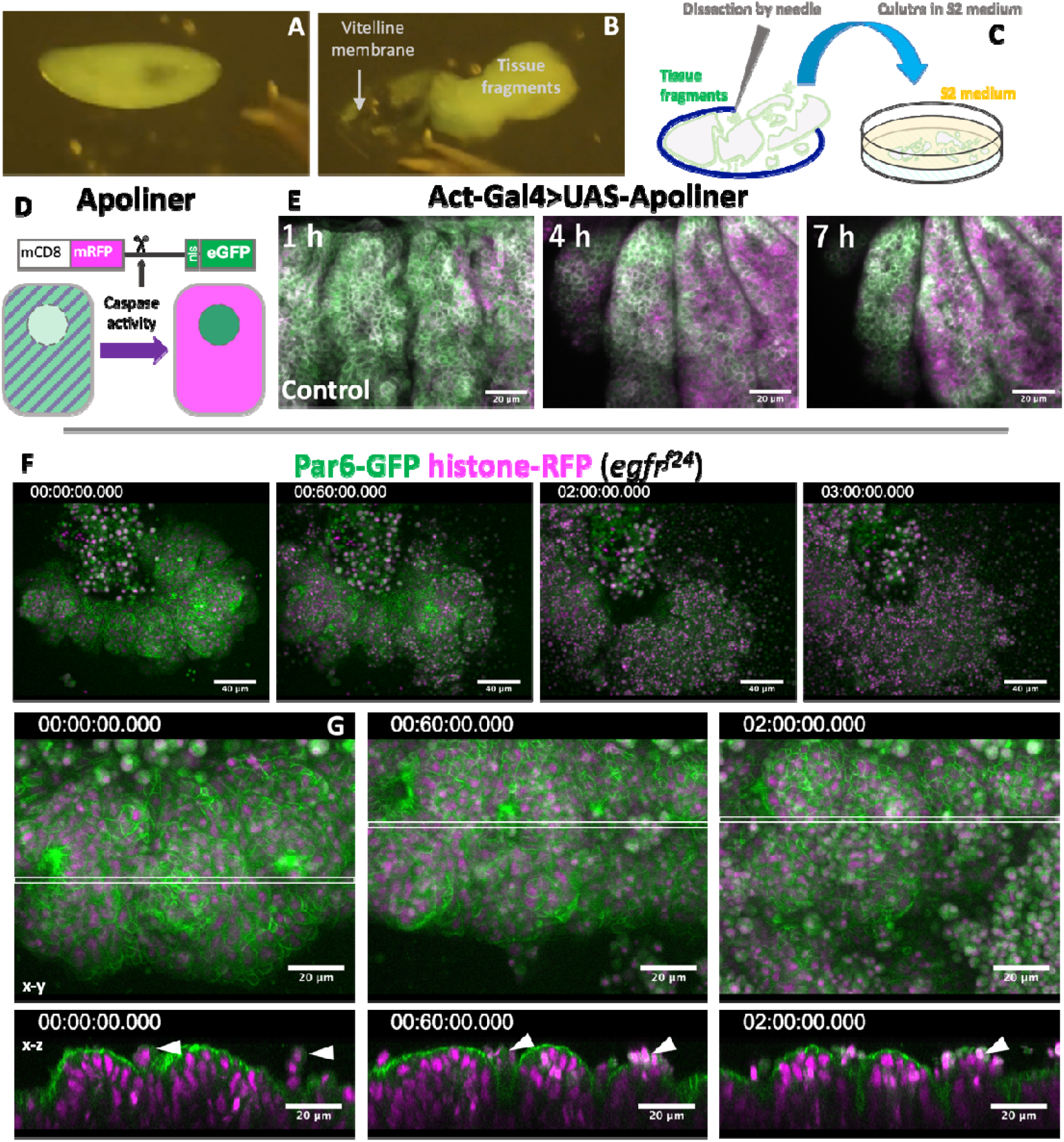
Rapid collapse of EGFR tissue fragments in explant cultures. **A-C**. The procedure of embryo dissection. **D**. Apoliner caspase activity detector. **E-G**. Time-lapse imaging of cultured tissue fragments. **E**. Apoliner activity in a control tissue fragment. **F**. EGFR mutant tissue fragment undergoing rapid disintegration. **G**. Early phase of EGFR mutant tissue culture, showing frequent apical cell extrusion (arrowhead). Imaging started within one hour after dissection.

### 2.6. EGFR signaling is required for protecting the epidermis from tissue wound

Next, we investigated whether EGFR protects tissues from injury. To this end, we used an approach to induce epithelial lesions with an infrared (IR) laser that reaches deep into the target and avoids the risk of damaging the vitelline membrane (Kato et al., 2016). Laser irradiation caused a wound (Figure. 6A, B; Miao and Hayashi, 2015) and induced the expression of di-phosphorylated ERK in 1–2 cell rows around the wounded region, as previously observed in tissue culture cells and *Drosophila* (Figure. 6C, Matsubayashi et al. 2004, Hiratsuka et al., 2015; Geiger et al., 2011; Mace et al., 2005; Wang et al., 2009). Under the same conditions, the ERK-FRET reporter was immediately activated in a region covering the adjacent segment and tracheal placode 5 min after wounding (Figure. 6D, 5 min). In the next 10 min, broad ERK activity was reduced to 1–2 cell rows at the wound edge, and this activity was sustained for at least 30 min after wounding (Figure. 6D, 10-15 min, Supplemental Movie. S6). Cables of myosin appeared at the wound edge 10 min after wounding (Figure. 6D’).

**Figure 6.**
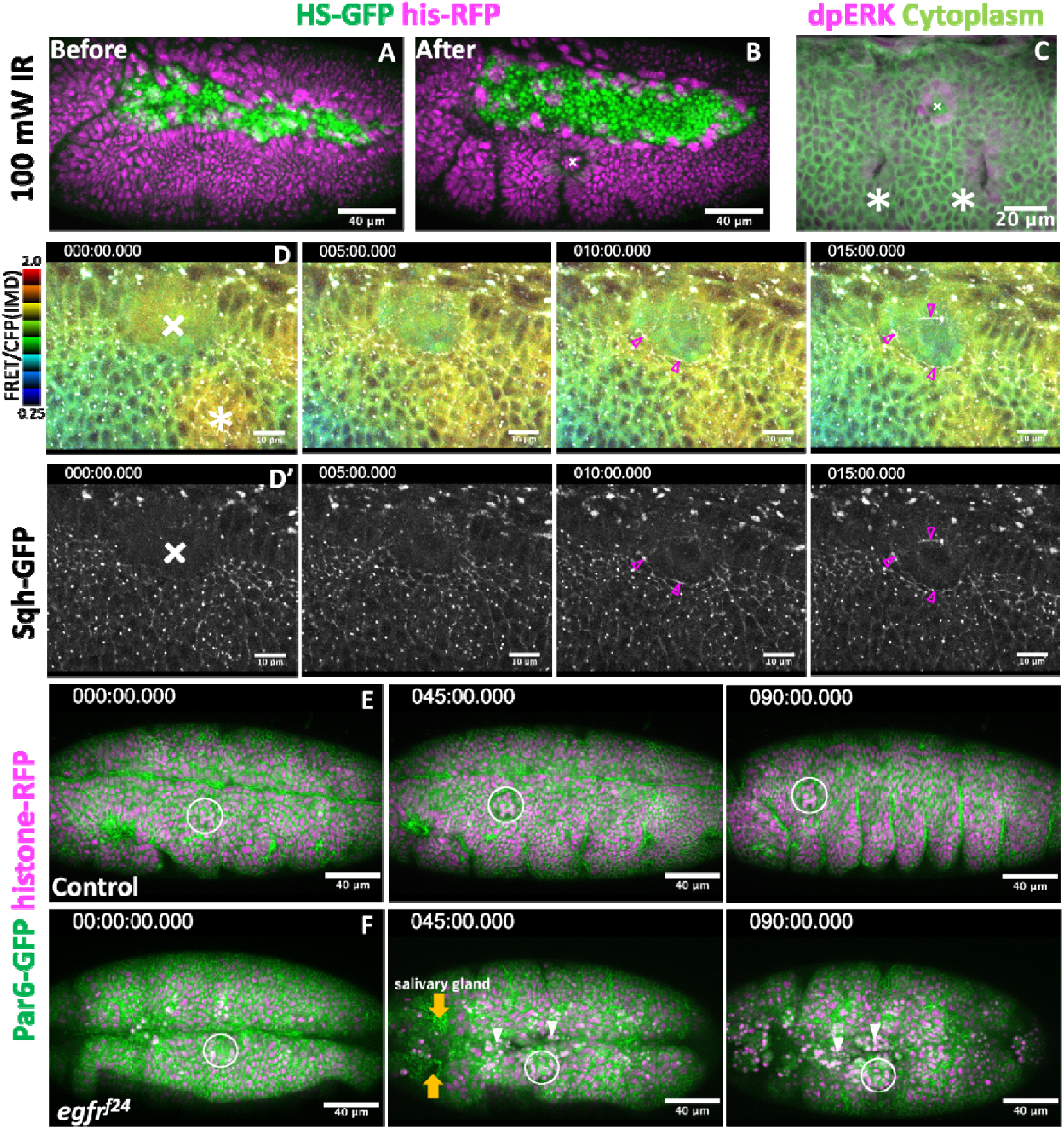
EGFR signaling is required for tissue repair. **A, B**. Tissue wounds treated with infrared laser illumination. **C**. ERK activation in the vicinity of the wound region (arrow). Cross marks; wounded site, Asterisks: tracheal placode. **D**. Simultaneous recording of ERK activity (FRET probe, upper row) and myosin activity (lower row). **E, F**. Laser wound damage is limited locally in the control group (**E**). Tissue disintegration spreads from the wound site in EGFR mutants (**F**). White circles indicate wounded site. Orange arrows indicate invagination of salivary gland.

Having established the conditions for laser wounding, we next applied this technique to EGFR mutants. A previous wounding experiment of stage 15 embryos demonstrated that wound healing was delayed in EGFR hypomorphic mutants (Geiger et al., 2011). Here, we chose to apply a wound in the region requiring EGFR (ventral epidermis) at the first thoracic segment in stage 11; the time and place at which we would not see any defect in EGFR mutants. Under these conditions, the damage introduced to control embryos was limited to the vicinity of the laser-illuminated area, and embryogenesis proceeded (N = 15 embryos, Figure. 6E, Supplemental Movie. S7). When the EGFR mutant was wounded, the damage expanded to the neighboring region, separate from the epithelial disintegration initiated from the head region, and the treated embryos collapsed ectopically in the thorax-abdominal regions (N = 10 embryos, Figure. 6F, Supplemental Movie. S7). Cross-sectional views showed that wounding caused ectopic detachment of the epithelia from the vitelline membrane around the vicinity of the wounding region (Supplemental Figure. 3A, B). In both control and EGFR-mutant embryos, ectopic detachment persisted for more than one hour (Supplemental Figure. 3C, D).

## 3. Discussion

In this study, we sought to understand the mechanisms that protect the epithelia from a variety of stresses present in developing embryos. By searching for conditions that exacerbate the apoptotic cell extrusion phenotype of EGFR mutants, we found that epithelial invagination severely enhances the phenotype. Invaginating epithelial cells create anisotropic contractile forces to induce the folding of the epithelia (Chung et al., 2017; Letizia et al., 2011; Nishimura et al., 2007; Sanchez-Corrales et al., 2018), which applies mechanical stress to the adherens junction. We found that EGFR-ERK signaling was upregulated at the site where the apical cell membrane detached from the vitelline membrane. We infer that the epithelium is mechanically supported by contact with the vitelline membrane and that their detachment activates EGFR-ERK signaling to suppress transient instability through elevated apical myosin activity. The functional overlap of the vitelline membrane and EGFR signaling was verified by the synergistic effect of removing both the vitelline membrane and EGFR signaling. Loss of EGFR function may reduce the threshold required for epithelial cells to tolerate instability from various sources. Here, we discuss the role of EGFR in stabilizing epithelia and how the vitelline membrane increases the robustness of morphogenetic movement.

### 3.1. EGFR promotes cell surface contractility and prevents apical cell extrusion

*Drosophila* EGFR has multifaceted roles, including the regulation of patterned gene expression, cell survival, and control of myosin contractility (Bergmann et al., 2002; Crossman et al., 2018; Hayashi and Ogura, 2020; Ogura et al., 2018; Price et al., 1989; Raz and Shilo, 1993; Schejter and Shilo, 1989). The latter two functions are relevant to this study. Myosin contractility and cell migration in cultured epithelial cells are controlled by the propagation of ERK activity through the activation of receptor tyrosine kinases (Aikin et al., 2020; Aoki et al., 2017), and the propagating wave of ERK has been observed in the mouse epidermis (Hiratsuka et al., 2015). In *Drosophila* epithelia, myosin accumulates at the adherens junction level to form cables spanning multiple cells, and functions to constrict junction length. Simultaneous imaging of myosin and ERK-FRET probes showed that ERK activation precedes myosin cable formation (Ogura et al., 2018, This work, Figure. 6). These myosin cables apply contractile forces to the apical side of the epithelium to promote tissue invagination (Kondo and Hayashi, 2013; Nishimura et al., 2007; Ogura et al., 2018). A recent study on *Drosophila* pupal notum reported that sporadic cell death triggers transient EGFR-ERK activity in the surrounding cells that promptly seal the space of dead cells and repair epithelial integrity (Valon et al., 2021). Concentrating myosin near the adherens junction ensures contractile tension is high in the apical surface so that the actomyosin cables function as a mechanical barrier to prevent cell extrusion to the apical side.

Rhomboids are the major activator of the EFGR ligand Spitz through intramembrane cleavage of the precursor form of Spitz in the Golgi apparatus (Shilo, 2016). The transcription patterns of *rhomboid*s closely correlated with the patterns of EGFR-dependent ERK activity. We showed that rhomboid is a major inducer of ERK activity in embryos. The appearance of sporadic apical cell extrusions in *rhomboid* mutant embryos indicated that the restriction of apical cell extrusions is controlled by *rhomboid*s. In EGFR mutants, a much larger number of cell deaths and tissue disintegration were observed, indicating that additional stimuli for EGFR must exist. It is unlikely that the other members of the seven rhomboid family genes fulfill that role because other members are either not expressed abundantly in the embryos or are non-proteolytic (Shilo, 2016). A potential alternative path for EGFR activation in the neighborhood of dying cells is stretch-induced EGFR activation, (Hino et al., 2020; Valon et al., 2021), in which apoptotic cells autonomously shrink and stretch forth to their neighbors and activate the EGFR-ERK pathway that stimulates contractile myosin cables in a pattern surrounding the dying cells. Such a myosin cable was observed, slightly after ERK activation, in the apical cell junction in the area of laser-induced cell killing (Figure. 7D), that undergoes basal ward movement and detachment from the vitelline membrane (Supplemental Figure. S3), suggesting that stretch-induced EGFR-ERK-myosin activation prevents apical cell extrusion in the embryo.

### 3.2. Vitelline membrane stabilizes epithelia

Apoptotic cells are extruded in either the apical or basal direction from cultured epithelium or zebrafish tissues (Eisenhoffer et al., 2012; Gu and Rosenblatt, 2012; Ohsawa et al., 2018; Tada, 2021). Basal cell extrusion is a common form of cell elimination in *Drosophila* imaginal discs (Ohsawa et al., 2018), and forced extrusion of polarity-deficient cells to the apical side causes luminal tumors (Vaughen and Igaki, 2016). In *Drosophila* embryonic epithelia, naturally or experimentally occurring apoptotic cells are mostly extruded into the basal side (Abrams et al., 1993, Figure. 2A, B). Although the basement membrane is a potential cue for basal ward cell extrusion in imaginal discs, the accumulation of basement membrane components is not complete until late-stage embryos (stage 16 or later) (Matsubayashi et al., 2017; Matsubayashi et al., 2020). This implies that another positional cue may restrict the direction of cell extrusion in the embryo. Close apposition of the apical cell membrane under pressure from the internal part of the embryo to the vitelline membrane serves as a physical barrier that restricts apical cell extrusion. In addition, part of the ectoderm adjacent to the hindgut invagination site was shown to be connected to the vitelline membrane via integrin adhesion (Münster et al., 2019). This adhesion system may offer a more robust stabilization mechanism.

### 3.3. Potential contribution of mitosis to tissue instability

We showed that clustered apoptosis accompanied the initial apical cell extrusion observed in EGFR mutants (Figure. 3). However, other clusters of apoptosis in the head and thoracic segments, which were tightly associated with the vitelline membrane, did not exhibit the apical cell extrusion phenotype (Figure. 4F, G). The only exception was the apoptotic cell cluster in salivary gland invaginations (Figure. 4F). This implies that a cluster of apoptosis is not sufficient to cause apical cell extrusion, and additional conditions, other than invagination, must cooperate with the EGFR mutation in the head region. We showed that clusters with high mitotic activity were present in the head (Supplementary Figure. S2). These mitotic cells were present in the region of clusters of apoptotic cells in EGFR mutants (Figure. 3C-F), whereas the apically extruded cells did not contain mitotic cells (Supplementary Figure. S2). Mitotic cells undergo cell rounding, in which actin and myosin are redistributed to the cell cortex to increase cortical tension (Kunda et al., 2008; Stewart et al., 2011) and maintain adhesion to their neighboring cells through the equatorial E-cadherin belt. Rounded mitotic cells bulge out of the apical surface to cause detachment from the vitellin membrane (Kondo and Hayashi, 2013, Supplemental Figure. S1C) and to apply stress to the weakened adherens cell junction, which is depleted of actin and myosin (Ragkousi and Gibson, 2014). We speculate that clustered mitotic activity in healthy cells in the head of EGFR mutants causes detachment from the vitelline membrane and enhances instability in the epithelia, leading to apical cell extrusion.

## 4. Concluding remarks

It should be noted that embryonic development in many experimental model organisms, such as mice, chicks, and zebrafish, can proceed outside of egg membranes, although the contribution of the mouse zona pellucida to embryo polarization has been documented (Kurotaki et al., 2007). Presumably, the relatively slow developmental time of these embryos (more than 3-fold longer than *Drosophila*) would reduce the negative impact of mitosis and invagination on epithelial stability and allow enough time for tissues to repair any damage that might occur during morphogenesis, making embryonic membranes dispensable. In addition, two cell adhesion systems, the septate junction mediating adhesion of lateral cell membranes and integrin-basement membrane adhesion systems, develop only in the late embryonic stages (stage 15 or later, Tepass and Hartenstein, 1994), leaving the task of maintaining epithelial cell adhesion during early morphogenesis mostly to the adherens junction. The simplicity of the adhesion system may allow flexibility in coping with rapid morphogenesis at the expense of an increased risk of tissue damage. The recruitment of the vitelline membrane as a supporting medium and EGFR signaling as a tissue repair mechanism permitted rapid and robust embryonic development of *Drosophila*, an advantage for adoption in the varying reproductive environment.

## 5. Materials and Methods

### 5.1 Key resource table

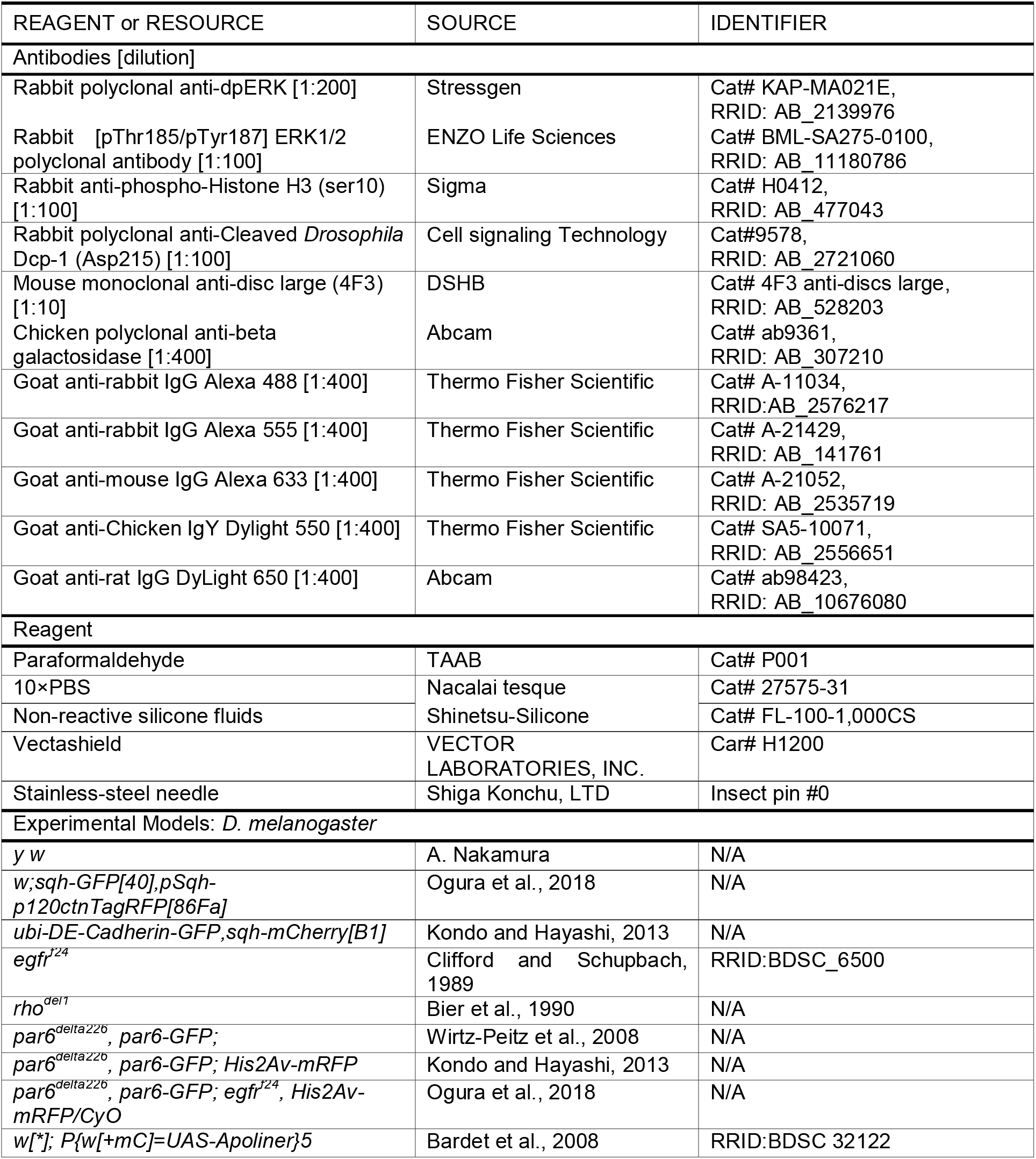
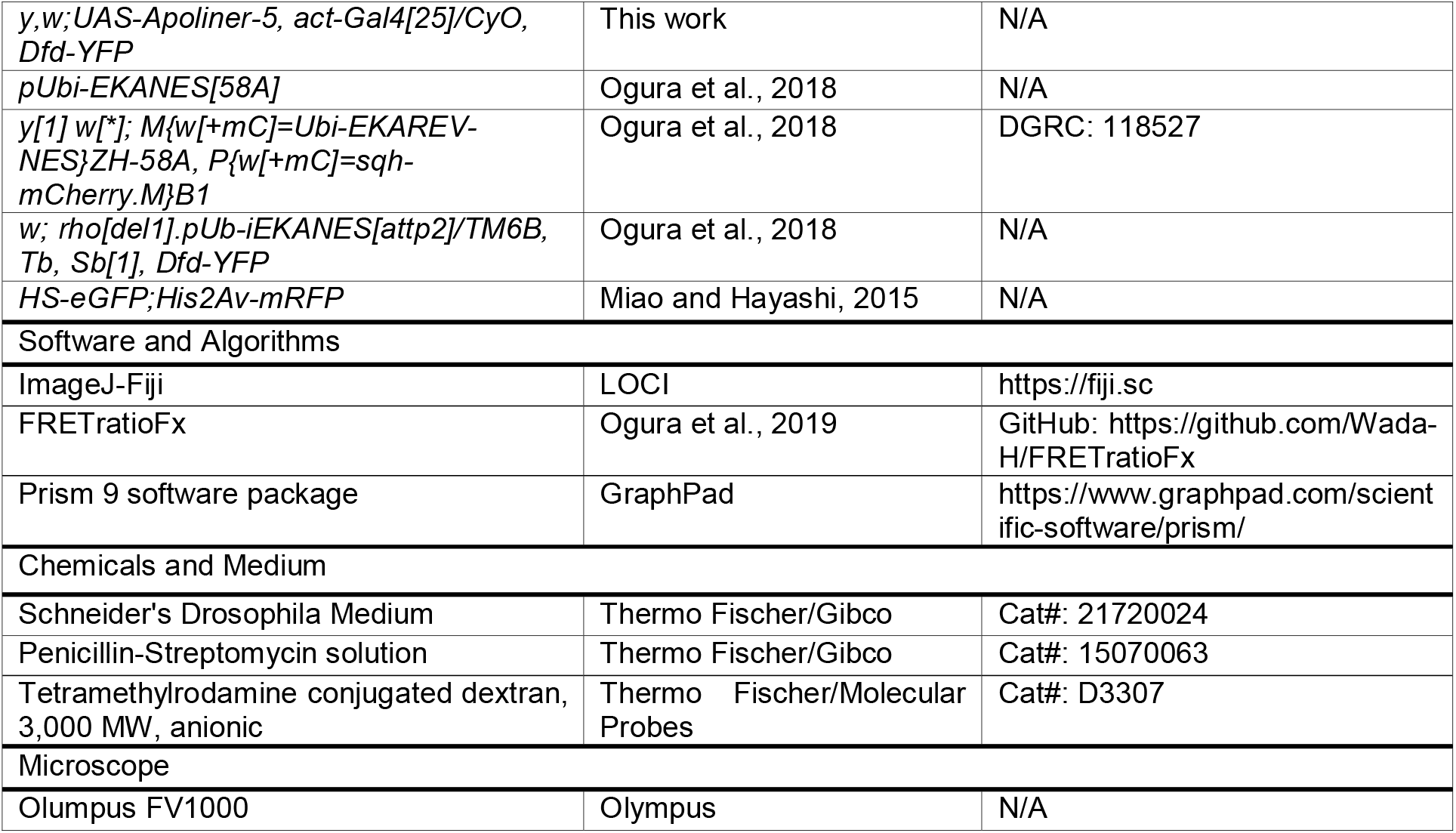

#### 5.2 Experimental Models

**All *Drosophila* strains were cultured in standard yeast-cornmeal media at 25°C. Embryos were staged according to morphological criteria and time course** (Campos-Orterga and Hartenstein, 1997).

### 5.3. Immunofluorescence and antibodies

Embryos were collected on apple juice-2 % agar plates covered with yeast. Chemically dechorionated embryos were fixed for 40 min with 4 % w/v paraformaldehyde in phosphate-buffered saline (PBS) and devitellinized by replacing the water phase with methanol with vigorous shaking. Immunostained embryos were mounted in Vectarshield and observed under a confocal microscope (Olympus FV1000) using UPlanSApo 20x NA0.75 with a 1.5 zoom, and UPLanSApO 60 x water NA 1.2. Each image was acquired as an 800 × 600-pixel image with an optically optimum z-interval.

### 5.4. Live imaging

Dechorionated *Drosophila* embryos were mounted on a glass-botto med dish (IWAKI) coated with glue extracted with heptane from a double-stick tape (Scotch). Each confocal image was acquired using a laser-scanning confocal microscope (FV1000, Olympus) with the following lenses: PlanApo N 60 x NA1.42 oil (most of the live imaging), UPlanSApo 60 x NA1.30 silicon (for FRET imaging with fluorescent dextran injection in Figure. 1F, G), UPlanSApo 30 x NA1.05 silicon (for observation of the whole embryo), and PlanApo 60 x NA1.40 oil IR lens (for IR-irradiation). Each image was acquired as a 640 × 480-pixel image with an optimum z-interval, every 3 or 5 min over 5 h with GaAsP detectors. The images were de-noised and projected using ImageJ-Fiji software. Förster resonance energy transfer imaging and data processing were performed as described previously (Ogura et al., 2018; Ogura et al., 2019).

### 5.5. Quantification of apical cell extrusion events

Progenies of *w, rho*^*del1*^, *pUbi-EKANES[attp2]/TM6B, Tb, Sb[1], and Dfd-YFP* stock were imaged for FRET activity using a multi-point time-lapse function. The genotype was determined by monitoring the expression of Dfd-YFP and the appearance of a fully mature embryo the next day. Only animals that completed embryogenesis were used for further image analysis. The x-y-z stack images of the lateral views of embryos were prepared from 4D time series, resized for the pixel size of 0.75 × 0.75 µm (x-y); x-z slice image series for each pixel of the y-axis were produced from the stacks and the number of slices which include at least one apically extruded cell was counted. The apical signal rate was defined as the frequency of slices that included apically extruded cells per total number of embryo slices. Statistical significance was evaluated by the Mann–Whitney U test using the Prism 9 software (GraphPad).

### 5.6. Fluorescence dextran injection

Dechorionated embryos in stages 11–12 were mounted on a cover glass (Matsunami) and covered with silicone oil (Shinetsu silicone). Solution of tetramethyl rhodamine-conjugated dextran, 3,000 MW (1 µg/µl) was diluted in PBS and injected into the perivitelline space of the embryos. The glass needles were prepared by pulling a glass capillary with a needle puller (Sutter).

### 5.7. Embryo dissection and culture

Eggs were collected overnight at 18 °C, and stage11–12 embryos were selected from the dechorionated embryos. They were placed on a polystyrene tissue culture dish (IWAKI) and covered with Schneider’s *Drosophila* medium supplemented with 10 % fetal bovine serum and antibiotics (penicillin and streptomycin). The vitelline membrane was punctured by pricking at the anterior end of the embryo with a stainless-steel needle (Shiga Konchu, LTD), and half of the disrupted embryos were pushed out of the vitelline membrane. Tissue fragments were collected without vitelline membrane from at least 10 embryos in each experiment and placed in poly L-Lysin-coated glass-bottomed dishes (Matsunami D141410) for confocal imaging. Epidermal tissue fragments facing the apical side toward the glass were imaged within 1 h after dissection.

### 5.8. Laser-induced wounding

The IR-LEGO system (IR-LEGO-1000, Sigma-Koki Co., Ltd., Saitama, Japan) was combined with a confocal microscope (FV1000, Olympus, Japan) equipped with GaAsP detectors. The IR laser was introduced through the lateral camera port of an inverted microscope (IX81; Olympus). The laser intensity was adjusted to 100 mW or 150mW to induce local wounds (Kamei et al., 2008; Kato et al., 2016; Miao and Hayashi, 2015). Time-lapse confocal images were captured immediately after IR laser application.

## Acknowledgments

We thank the Kyoto *Drosophila* Stock Center, Bloomington Stock Center, and Developmental Studies Hybridoma Bank (DSHB) for providing fly stocks and antibodies. We thank Yosuke Ogura and Housei Wada for their expert technical assistance and members of the Hayashi lab for their comments on the manuscript. K.Y. is the RIKEN Junior Research Associate. This study was supported by a Grant-in-Aid for Scientific Research [19H00996 to S.H.] from MEXT, Japan.

## Supplementary Figures

**Supplementary Figure S1.**
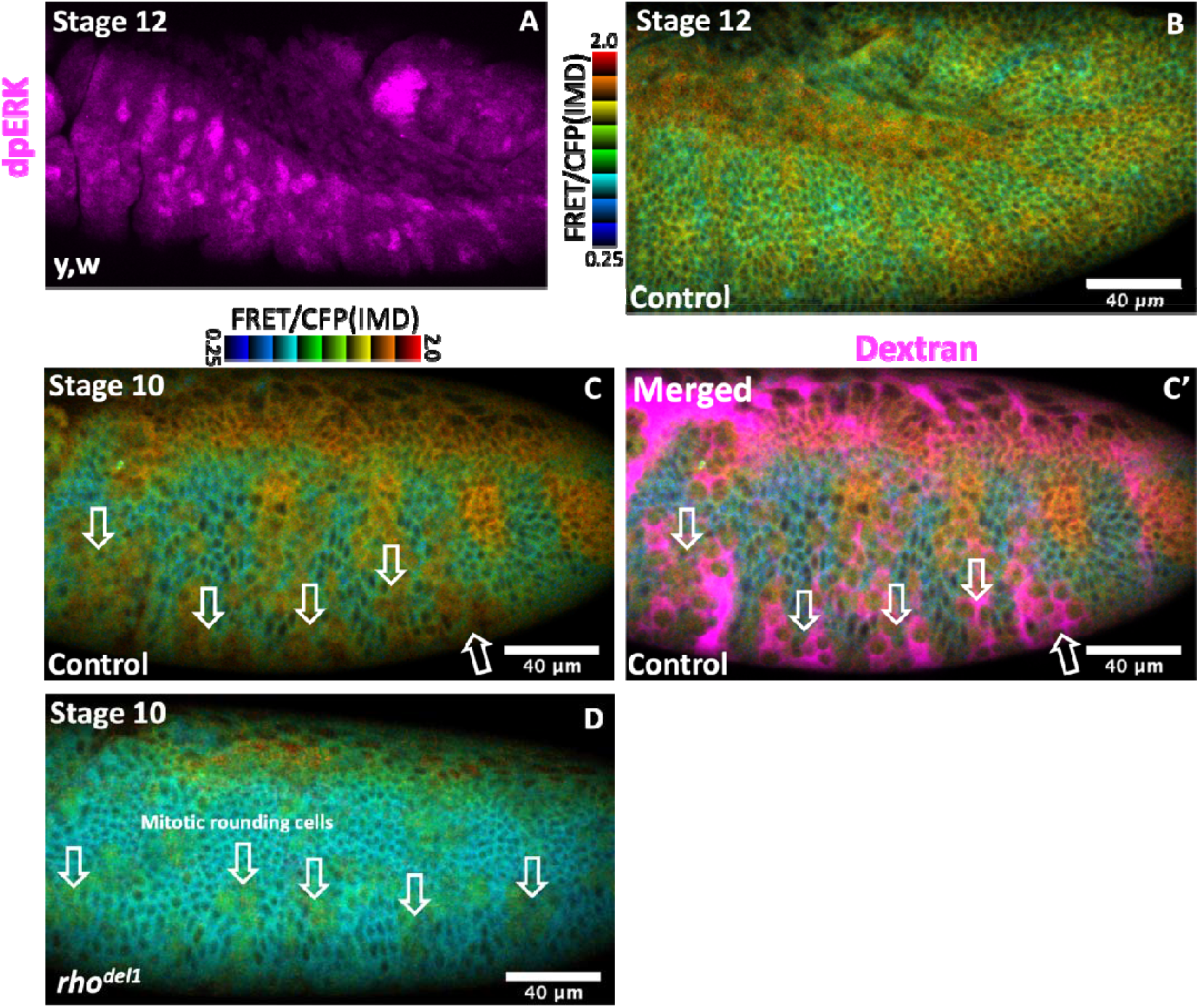
**A, B**. dpERK staining. (**A**) ERK-FRET images of stage 12 embryos. **C, C’**. ERK-FRET image (**C**) and an overlay of perivitelline dye (**C’**) in stage 10 embryos. Arrows indicate FRET high cells with an open perivitelline space. These are mitotic cell clusters that detach from the vitelline membrane. The FRET signal reflects the CDK1 signal and persists in *rho* null mutant embryos (**D**).

**Supplementary Figure S2.**
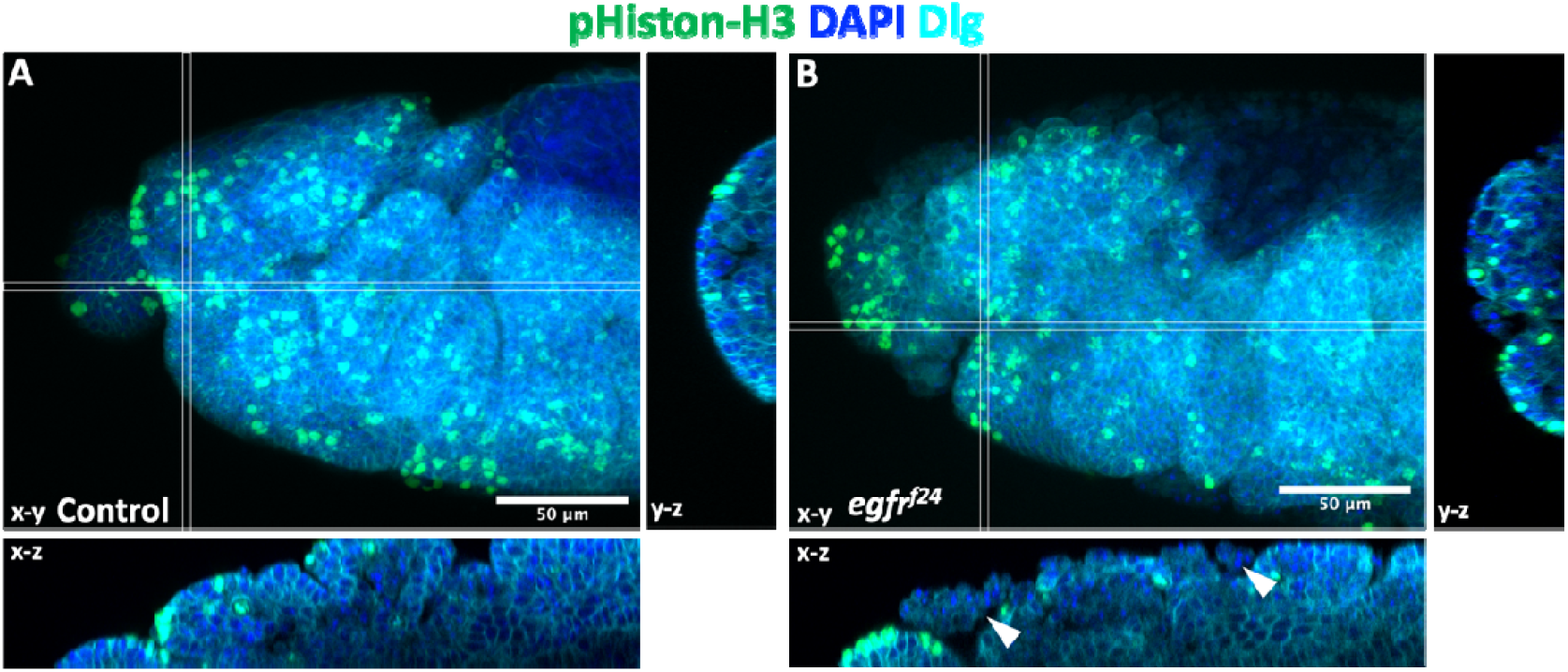
Patterns of mitotic cells in the head. Mitotic cell marker phosphorylated histone H3 staining of control (**A**) and EGFR mutant (**B**) cells in stage 12. Closed arrowhead indicate the cluster of apically extruded cells.

**Supplementary Figure S3.**
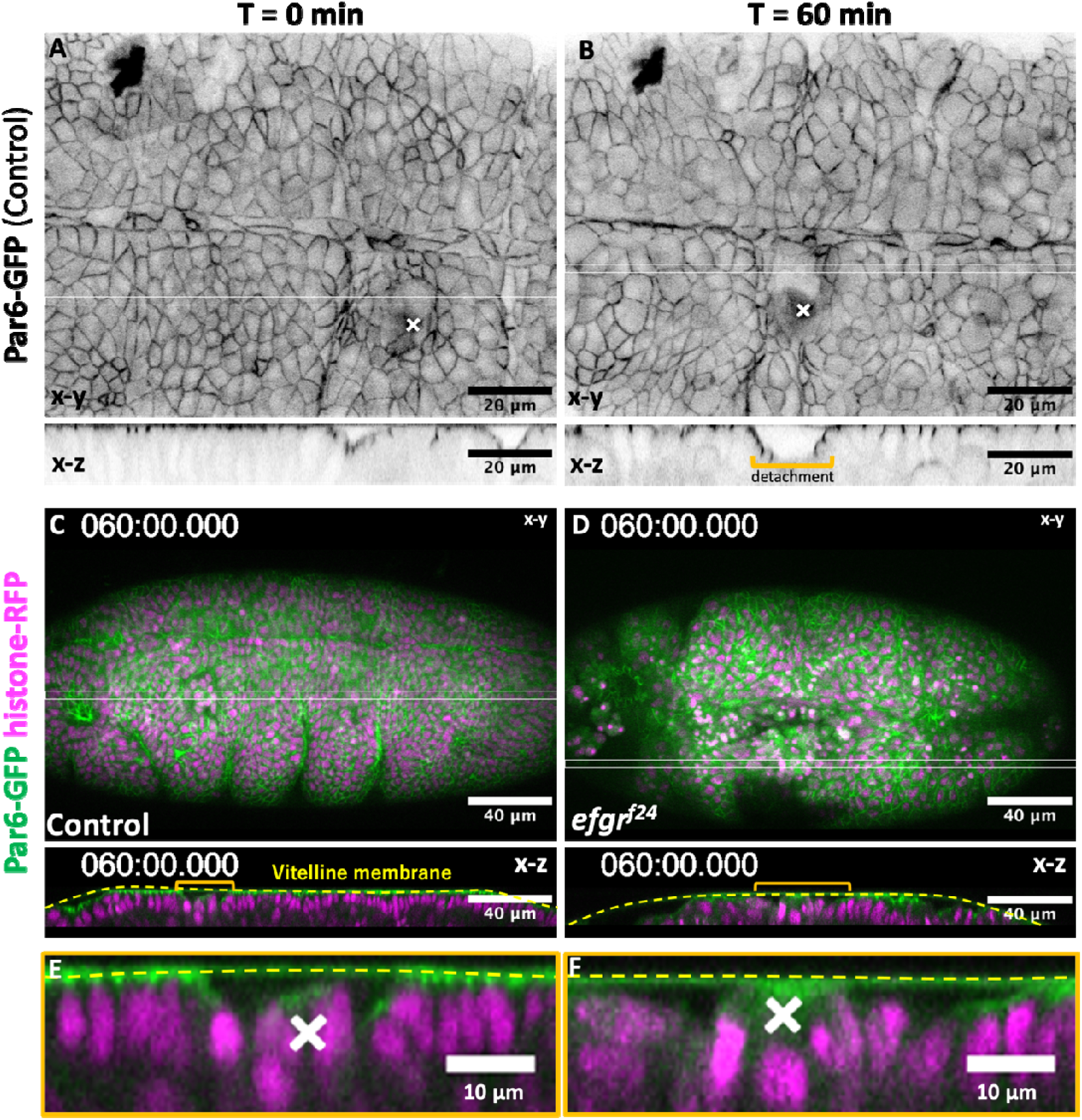
Wounding causes detachment of epithelia from the vitelline membrane. **A, B**. X-Y and X-Z sections of the region near the wound site. The x-z section of the double horizontal line is shown. **A**. Immediately after laser wounding. The cells were attached to the vitelline membrane. **B**. 60 min later, a large gap appeared. The gap was observed 5 min after wounding and gradually increased in size. **C-F**. Cross-sectional views of wounded regions of the control (**C, E**) and EGFR mutant (**D, F**) 60 min after wounding. Note in enlarged views (**E, F**) that apical Par6-GFP was maintained in the control (**E**), but lost in the mutant (**F**). Related to Figures 6 E, F, and Supplemental movie S7. Cross marks; wounded site.

## Supplemental Movies

**Supplementary Movie S1**. ERK-FRET imaging of embryos injected with a fluorescent tracer (rhodamine dextran) in the perivitelline space. Related to Figure 1F, G.

**Supplementary Movie S2**. Head phenotype of the EGFR mutant. Imaging was started in the late stage 10. Time t = 0 was set at the time of full opening of the tracheal pit at T2. Related to Figure 2D, E.

**Supplementary Movie S3**. Apical cell extrusion in the EGFR mutant head. The time to the first apically extruded cell was set as t = 0. This cell was tracked by adjusting the positions of the x-z and y-z slices. Related to Figure 3G.

**Supplementary Movie S4**. Apical cell extrusion at the salivary gland invagination site of EGFR mutants. Ventral views of the head and thoracic regions of the two embryos were obtained. Time t = 0 was set at the beginning of invagination. Related to Figure 4F, G.

**Supplementary Movie S5**. Tissue explant experiment. Control and EGFR mutant tissue fragments, removed from the vitelline membrane, and cultured *in vitro*, were imaged. Related to Figure 5G.

**Supplementary Movie S6**. Responses of myosin (myosin light chain GFP) and ERK signaling (ERK-FRET) to laser-induced wounding in control embryos. Related to Figure 6D.

**Supplementary Movie S7**. Laser-induced wounding in EGFR-mutant embryos. Embryos at stage 12 prior to tissue collapse in the thorax were chosen for the experiment. Related to Figure 6E, F.

